# Coarse-grained chromatin dynamics by tracking multiple similarly labeled gene loci

**DOI:** 10.1101/2025.02.27.640402

**Authors:** Alexander Mader, Andrew I. Rodriguez, Tianyu Yuan, Ivan Surovtsev, Megan C. King, Simon G. J. Mochrie

**Affiliations:** Department of Physics, Yale University, New Haven, Connecticut 06511, USA; Integrated Graduate Program in Physical and Engineering Biology, Yale University, New Haven, Connecticut 06511, USA; Department of Molecular Biophysics and Biochemistry, Yale University, New Haven, Connecticut 06511, USA; Department of Cell Biology, Yale School of Medicine, New Haven, Connecticut 06520, USA; Department of Molecular, Cellular, and Developmental Biology, Yale University, New Haven, Connecticut 06511, USA; Department of Applied Physics, Yale University, New Haven, Connecticut 06511, USA

## Abstract

The “holy grail” of chromatin research would be to follow the chromatin configuration in individual live cells over time. One way to achieve this goal would be to track the positions of multiple loci arranged along the chromatin polymer with fluorescent labels. Use of distinguishable labels would define each locus uniquely in a microscopic image but would restrict the number of loci that could be observed simultaneously, because of experimental limits to the number of distinguishable labels. Use of the same label for all loci circumvents this limitation but requires a (currently lacking) framework for how to establish each observed locus identity, i.e. to which genomic position it corresponds. Here we analyze theoretically, using simulations of Rouse-model polymers, how single-particle-tracking of multiple identically-labeled loci enables determination of loci identity. We show that the probability of correctly assigning observed loci to genomic positions converges exponentially to unity as the number of observed loci configurations increases. The convergence rate depends only weakly on the number of labeled loci, so that even large numbers of loci can be identified with high fidelity by tracking them across about 8 independent chromatin configurations. In the case of two distinct labels that alternate along the chromatin polymer, we find that the probability of the correct assignment converges faster than for same-labeled loci, requiring observation of fewer independent chromatin configurations to establish loci identities. Finally, for a modified Rouse-model polymer, that realizes a population of dynamic loops, we find that the success probability also converges to unity exponentially as the number of observed loci configurations increases, albeit slightly more slowly than for a classical Rouse model polymer. Altogether, these results establish particle tracking of multiple identically- or alternately-labeled loci over time as a feasible way to infer temporal dynamics of the coarse-grained configuration of the chromatin polymer in individual living cells.

**SIGNIFICANCE:** In spite of recent success in elucidating its spatial organization, chromatin’s time-dependent, dynamical behavior remains far less studied, and correspondingly much less understood. To address the critical need to elucidate chromatin dynamics, this paper proffers a route towards an experimental characterization of coarse-grained chromosomal dynamics, via particle tracking of multiple labeled loci, labeled with just one or two different fluophor colors or intensities. Theoretically, we show that particle tracking of multiple identically labeled loci across only about 8 independent chromatin configurations should be a feasible way to establish the time-dependent, coarse-grained configuration of the chromatin polymer in individual living cells.

## INTRODUCTION

The 4D nucleome project (1) has been phenomenally successful in advancing our understanding of chromatin’s spatial organization. Preeminently, crosslinking- and sequencing-based chromatin-configuration-capture experiments (Hi-C, *etc*.) now can measure the population-averaged relative contact probability between all pairs of genomic loci, typically presented as a map of contact probabilities (2–11). Hi-C measurements have inspired theoretical models of chromatin organization that can reproduce measured Hi-C maps, based on three coexisting mechanisms, operating at different genomic scales. First, on several-megabase scales, “checkerboard patterns” in Hi-C maps, encompassing distant contacts on the same chromosome, as well as contacts on different chromosomes, have led to a block co-polymer-inspired picture of chromatin compartments, comprised of several types of epigenetically-distinguished heterochromatin and euchromatin (12–25). Second, on the tens-to hundreds-of-kilobase scales, overlapping squares of high contact probability on the same chromosome, termed topologically associated domains (TADs), have led to the loop extrusion model in which a population of loop extrusion factors, identified as cohesin complexes, establishes a dynamic steady-state of preferentially-located chromatin loops (26–35). Third, at few-kilobase scales, high-resolution Hi-C reveals patterns of contacts that arise from nucleosomal structures (36–38). In addition to Hi-C, super-resolution microscopy, combined with sequential labeling of numerous target sequences via fluorescence in-situ hybridization (FISH), has directly observed 3D chromatin conformations in individual fixed cells (39–42), revealing high variability from cell to cell, consistent with the stochasticity implied by loop extrusion and block-copolymer models.

However, both Hi-C and FISH suffer from the same essential restriction, namely that they use fixed cell preparations, and hence provide limited information concerning the fourth dimension, time, *i*.*e*. concerning chromatin dynamics. On the other hand, changes in chromatin conformations underlie numerous nuclear and genomic processes, including regulation of gene expression, DNA repair, chromosome segregation, *etc*.. To properly understand these processes and fully realize the goals of the 4D nucleome project, it is paramount to be able to monitor how chromatin configurations evolve in time, for which we must look beyond Hi-C and FISH.

One successful and widely applied approach for probing the dynamics of an individual gene locus in living cells involves engineering an insertion near that locus of tandem arrays of a target DNA sequence, while also incorporating a fusion comprising a DNA-binding protein, that specifically binds the target sequence, fused to a fluorescent protein. For example, tandem arrays of the *lacO* sequence, inserted near the locus of interest, can be fluorescently-labeled by LacI-GFP, which binds *lacO*, allowing the motions of the so-labeled locus to be tracked versus time via GFP-based fluorescence microscopy (43–47).

To monitor the dynamics of multiple known loci simultaneously, it seems natural to label each locus with a uniquely-colored fluorophor, so that color establishes the genetic identity of each locus unambiguously. Indeed, a number of studies have examined the relative motion of two differently fluorescently-labeled loci in live cells (48–54). However, simultaneously labeling multiple loci, each with a different color, is a challenging genetic engineering task that would require the creation of cells that incorporate multiple tandem arrays of different target sequences, as well as all of the corresponding DNA binding protein-fluorescent protein fusions. Moreover, the number of different flurophors that can be spectrally-resolved is limited by the relatively broad emission spectra of available fluorophors. For these reasons, an approach that relies on labeling with multiple unique colors alone seems impractical for more than a few loci. So, what can we do?

Within the context of simple polymer models, genomic proximity along the chromatin polymer leads to spatial proximity on average. Specifically, the time-averaged square separation between genomically closer loci is less than that between genomically distant loci. To the extent that chromatin can be described by such models, it follows that by time-averaging particle tracking data, it should possible to unambiguously identify loci that are genomic neighbors, from which it is straightforward to deduce the ordering of the loci along the chromatin polymer. Knowing the ordering of loci along the chromatin polymer establishes the assignment of loci to their known genomic positions to within a complete reversal of the ordering. Experimentally, this issue can be resolved by studying cells that include a distinguishable, fluorescent nuclear landmark that is known to be proximal to a specific region of the chromatin under study. For example, in fission yeast the chromosome centromeres are proximal to the spindle pole body, which could be fluorescently labeled. Alternatively, one of the “end” labels could be at a genomic distance recognizably different from the rest of the inter-loci distances, thus establishing directionality.

Although tracking multiple loci over time is guaranteed to yield the correct ordering of loci along a polymer in the limit of an infinitely long measurement, for this method to be practically useful, it must yield the correct ordering of fluorescently-labeled loci with high probability within the duration of a fluorescent microscopy experiment. Unfortunately, under fluorescence excitation, fluorescent proteins photobleach, after which they no longer fluoresce. As a result, bleaching limits the duration over which each locus can be tracked. Therefore, if particle tracking is to be a practical route to loci identification, it must establish loci identity faster than the loci bleach. The goal of this paper is to examine theoretically via simulations what minimum configurational sampling during loci tracking measurements is necessary to accurately determine the correct ordering of multiple fluorescently labeled loci along the chromatin polymer.

Complementary to the present work is Ref. (55), whose focus is on the probability that multiple similarly-labeled loci are faithfully tracked from one microscopy image to the next. Ref. (55) establishes criteria, involving the time step between successive images, the genomic separation of labeled loci, *etc*., for achieving high-fidelity tracking. In this paper, we take perfect tracking for granted.

The remainder of this paper is organized as follows. Because we model the dynamics of the chromatin as a Rouse polymer, following many previous researchers (49, 56–58), in *Materials and Methods*, we first briefly review how we carry out simulations(35, 47). Then, we describe our procedure for determining loci identity from simultaneous multiple loci tracking and how we calculate the success probability that the procedure returns correct loci assignment. In *Results*, we first present results for the case when all loci are identically-labeled and show that the success probability depends (1) on the number of observations of loci configurations, (2) on the time step between the observations, and (3) on the genomic separation between labeled loci. The success probability converges exponentially to unity with increasing number of observations. Next in *Results*, we consider the case of two distinguishable loci labels (by color or intensity) that alternate along the chromatin polymer. Then, we examine the case of same-label loci for a Rouse polymer with dynamic loops generated according to the version of the loop extrusion model, described in detail in (59). In both cases, we again find that the success probability converges exponentially to unity with increasing numbers of observations. For the polymer with loops and identically-labeled loci, the convergence rate is slightly slower relative to the classical Rouse polymer. Unsurprisingly, the alternating labeling scheme leads to a convergence rate that is approximately two-fold faster than the single labeling scheme. Remarkably, for all cases considered, the convergence rate depends only weakly on the number of labelled loci, so that even multiple similarly labeled loci can be identified/assigned to the correct genomic location with high fidelity by tracking them across a limited number of independent chromatin configurations, even though increasing the number of same-label loci leads to a combinatorial growth of possible loci assignments. Finally, in *Discussion*, we conclude: Particle tracking of multiple identically or alternatingly labeled loci across a limited number of independent chromatin configurations, even considering experimental limitations such as photobleaching and successful tracking, is a feasible route to establishing the time-dependent, coarse-grained configurations of the chromatin polymer in individual living cells.

## MATERIALS AND METHODS

### Rouse-model polymer simulations

#### Polymers without loops

We carried out simulations of Rouse model polymers with free ends, as described in detail in Refs. (35, 47). In brief, we first simulate a time series of each Rouse-model normal coordinate, *e*.*g*. along the *x*-direction, *X*_*p*_ (*t*), via (60):

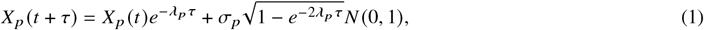

with

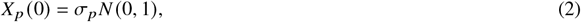

where *λ* _*p*_ is the *p*-th eigenvalue of the Rouse model dynamical matrix, 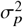 is the variance of *X*_*p*_, and *N* (0, 1) is a zero-mean Gaussian random variable with standard deviation equal to 1.

The position of bead *n* along the polymer, *x*_*n*_ (*t*) with *n* = 1, 2, … *M*, is then found via

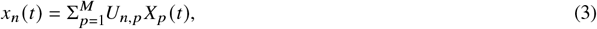

where *U*_*n*,*p*_ is the transformation matrix from normal mode coordinates to bead coordinates. (We ignore the *p* = 0 mode because it corresponds to the polymer’s center of mass motion and does not contribute to the separations between beads.) Analogous results also apply along the *y*- and *z*-directions, yielding time series for *y*_*n*_ (*t*) and *z*_*n*_ (*t*). For the classical Rouse model,

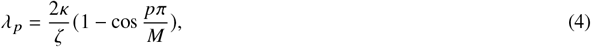

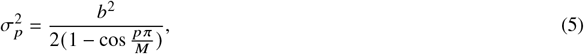

and

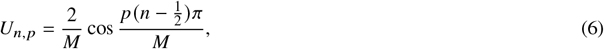

where *M* is the total number of Rouse-model beads, *κ* is the stiffness of the springs between Rouse-model beads, *ζ* is the beads’ friction coefficient, and 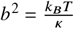 is the mean square separation along the *x*-axis between neighboring beads. We set the length and time units such that *b*^2^ = 1 and 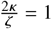.

The top panel of Fig. 1 illustrates one configuration of a classical Rouse polymer comprising 101 beads, connected by 100 springs. To model labeled loci, we identify a subset of the {(*x*_*n*_ (*t*), *y*_*n*_ (*t*), *z*_*n*_ (*t*))} as the positions of the labeled loci. For example, the second panel from the top of Fig. 1 shows the same polymer, but additionally highlights as the green circles, nine labeled beads at *n* = 10, 20, 30, 40, 50, 60, 70, 80, and 90. The third panel from the top shows these labeled beads without the connecting polymer. We seek to infer the correct assignment of the labeled beads along the polymer from their time-averaged separations. The bottom panel of Fig. 1 shows the same polymer as in the upper three panels, with both the positions of the nine labeled loci, and the coarse-grained reconstruction of the polymer configuration, obtained by connecting the labeled loci in the correct order along the chromatin.

**Figure 1.**
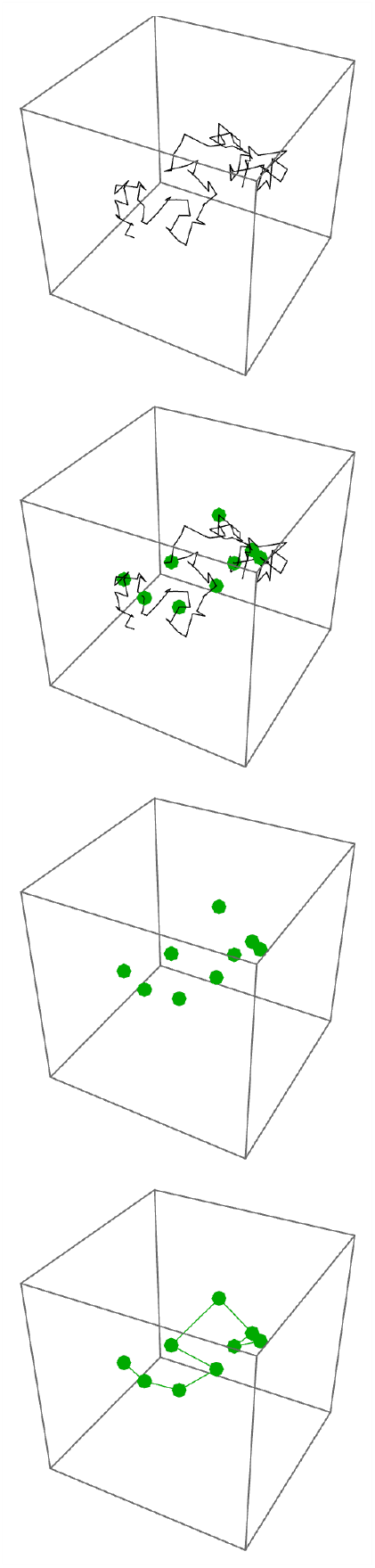
Top: The black line illustrates the configuration of a 101-bead Rouse-model polymer at one instant in time. Second from top: The same polymer as in the top panel, but now including the positions of nine labeled loci at beads 10, 20, 30, 40, 50, 60, 70, 80, and 90. Third from top: The same polymer as in the upper two panels, but now illustrating only the positions of the nine labeled loci. Bottom: The same polymer as in the upper three panels, with both the positions of the nine labeled loci, and the coarse-grained reconstruction of the polymer configuration, obtained by connecting these labeled loci in the correct order along the chromatin.

#### Polymers with loops

The canonical Rouse model polymer has long been used to describe the dynamics of chromatin (49, 56–58). However, Hi-C contact maps in conjunction with loop-extrusion-based models suggest that chromatin realizes a dynamic population of chromatin loops, inhomogeneously and stochastically distributed along the chromatin polymer. To investigate the role of chromatin loops, we carried out simulations of a modified Rouse model (35, 47), that includes loops. Specifically, we first carried out a Gillespie-type simulation of the loop configurations of the region from 0.2 Mb to 1.4 Mb of fission yeast’s chromosome 2, for 10^7^ loop extrusion events, which is sufficient to fully sample the ensemble of possible loop configurations, using the version of the loop extrusion model, described in detail in Ref. (59). Next, we randomly selected 10^5^ of the resultant loop configurations without regard to their original ordering. For each of these random loop configurations, we then generated a random Rouse polymer configuration, as described in detail in (35). Thus, in this case only, the averaging involves looped polymer configurations, that are completely independent of each other. For these looped-polymer simulations, we represented chromatin regions comprising 1.2 × 10^6^ base pairs via 4800 Rouse beads, so that the mean loop size of 22 kb base pairs corresponds to 88 beads (59).

### Measuring success probability

The polymeric nature of the chromatin dictates that the time-averaged separation between genomically close loci is necessarily less than that between genomically more distant loci. This suggests the following strategy for mapping observed loci to their correct genomic positions:

1. Track the positions of all loci over time. Here we assume that the tracking is “perfect”, i.e. each trajectory reflects motion of one locus, without changing its identity.
2. Arbitrarily assign genomic loci to each trajectory and calculate the cumulative sum of all neighboring loci square separations over all time points.
3. Repeat step 2 for all possible permutations of how to assign loci to genomic position.
4. Select a loci-to-trajectory assignment with the smallest cumulative square separation between neighboring loci as a “correct” mapping of observed loci to genetic positions.
5. Because the correct assignment of loci to genomic positions, *i*.*e*. the “ground truth”, is known in our simulations, by repeating this procedure for multiple simulated polymer configurations, and counting how many times the assignment of loci was indeed correct, we calculate the “success probability” that the procedure yields the correct assignment for a new set of polymer configurations.

Since the number of assignment permutations grows factorially with the number of labeled loci, *n*, it is computationally most efficient to use an algorithm for solving the traveling salesperson problem (TSP) to find the loci-to-trajectory assignment that minimizes the cumulative square separation between neighboring loci. This TSP-based approach has complexity *O* (*n*^2^2^*n*^) rather than the complexity *O n*! of the direct enumeration approach. However, for consistency throughout this paper, we employed the direct enumeration approach. In a number of cases, we verified that a TSP algorithm produced results identical to direct evaluation.

## RESULTS

### Success probability based on a single polymer configuration

We started by evaluating the success probability of mapping labeled loci to their correct genomic positions (i.e. to their polymer chain coordinates) based on a single observation, namely *P*_1_, by selecting the mapping that minimizes the total square separation between all pairs of neighboring labeled loci for a single polymer configuration. Fig. 2 display the values of *P*_1_ as a function of the number of labelled loci for a classical Rouse model polymer in the case that all loci are labelled identically (green circles) and the case that loci are alternatingly labeled with two distinguishable labels (blue circles). We also considered the case of a Rouse polymer with dynamic loops with identically labelled loci (red circles). The success probability for two identically-labeled loci or three alternatingly-labeled loci equals one because only one configuration of neighbors is possible in these cases. However, as the number, *n*, of labeled loci (and, hence, the number of possible mapping combinations) increases, the success probability decays rapidly. In each case, this behavior can be well described by exponential functions, shown as the lines in Fig. 2. which correspond to

**Figure 2.**
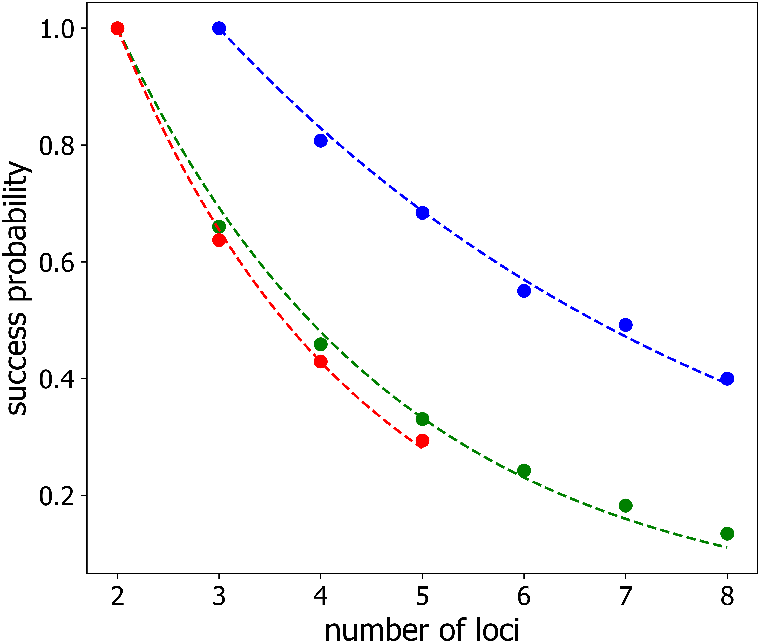
Success probability *P*_1_ to assign loci to their genomic positions correctly from a single observation (as outlined in *Matherials and Methods*) versus the number of labeled loci: for identically-labeled loci (*green*), for alternatingly-labeled loci (*blue*), and for identically-labeled loci on a polymer with loops (*red*). The lines correspond to single exponential fits (Eq. 7 or Eq. 10) to the data .

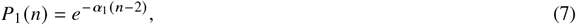

in the case of identically-labeled loci, and

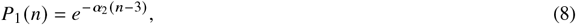

in the case of alternatingly-labeled loci (blue line). The best-fit values of *α*_1_ are *α*_1_ = 0.367 ± 0.010 in the case of the classical Rouse model (green line), and *α*_1_ = 0.424 ± 0.009 in the case of the Rouse model with loops (red line). The best fit value of *α*_2_ is *α*_2_ = 0.188 ± 0.005, corresponding to the blue line, so that the success probability decreases with increasing number of labeled loci approximately two times more slowly for alternatingly-labeled loci than for identically-labeled loci.

The rapid decay of successful mapping, based on a single observation, reflects the fact that the number of possible configurations is growing combinatorially with the number of labeled loci and that loci move stochastically. The stochastic character of the polymer configuration means that, while the distance between neighboring loci is smaller than the distance between more distant loci in an infinite-time average, this circumstance may not hold at any given moment of observation.

### Success probability for identically labeled loci tracked over time

As discussed above, however, we should expect that repeated sampling, while perfectly tracking labeled loci, will overcome polymer configuration stochasticity and increase the probability of successful mapping. Accordingly we next examined how the number of configurations sampled affects the probability of correct mapping. In this case, we considered a polymer configuration with *n* identically- and equidistantly-labeled loci evolving in time (governed by Eq. 1 as detailed in *Materials and Methods*). We sampled the polymer configuration *m* times, separated one from the next by a time step, *τ*. Then, we mapped the observed spatial loci positions to the corresponding genomic positions based on the minimum summed square-separation between loci, averaged over *m* polymer configurations, as described in *Materials and Methods*.

Fig. 3 shows the “success probability”, *P* (*m*) , that this procedure correctly predicts the loci assignments, plotted as a function of the number of configurations sampled/averaged, *m*, for polymers with *n* = 3, 4, 5, and 6 labeled loci (black, red, blue and green points, respectively), separated along the chain by 3 and 6 unlabeled loci (squares and triangles, respectively) for a sampling time step of *τ* = 1. As reasoned above, increasing the sampling indeed improves the probability of correctly mapping loci, with *P* (*m*) approaching an asymptotic value of 1 for large values of *m*. The solid lines in Fig. 4 are the best-fits to a simple exponential form:

**Figure 3.**
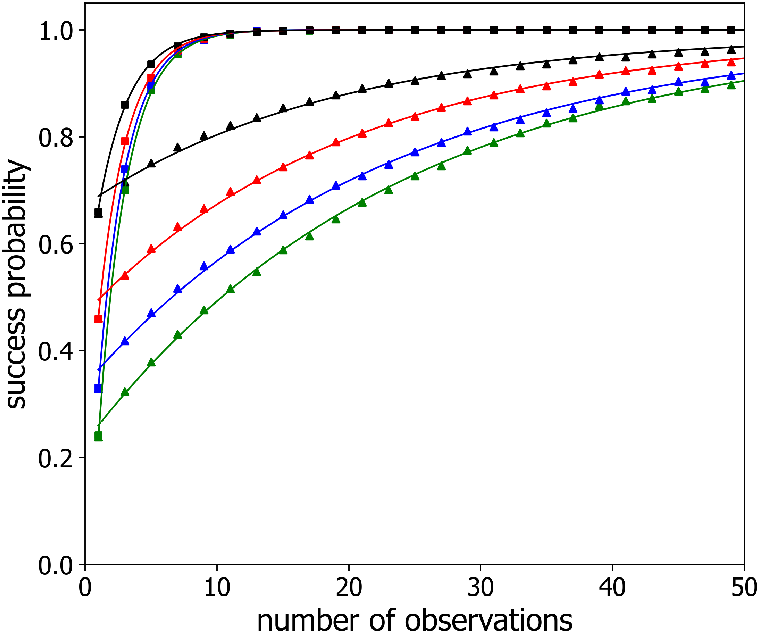
Success probability *P* (*m*) versus the number of observations *m* for observation time steps *τ* = 1 (*triangles*) and *τ* = 1000 (*squares*) and different numbers of labeled loci *n*: *n* = 3 (*black*), *n* = 4 (*red*), *n* = 5 (*blue*), *n* = 6 (*green*). Simulated Rouse polymer was comprised of 70 beads, and labeled beads/loci were located near the center with locus-to-locus separation of four beads. For clarity, every other data point is plotted. The solid lines are single exponential fits (Eq. 9) to the data.

**Figure 4.**
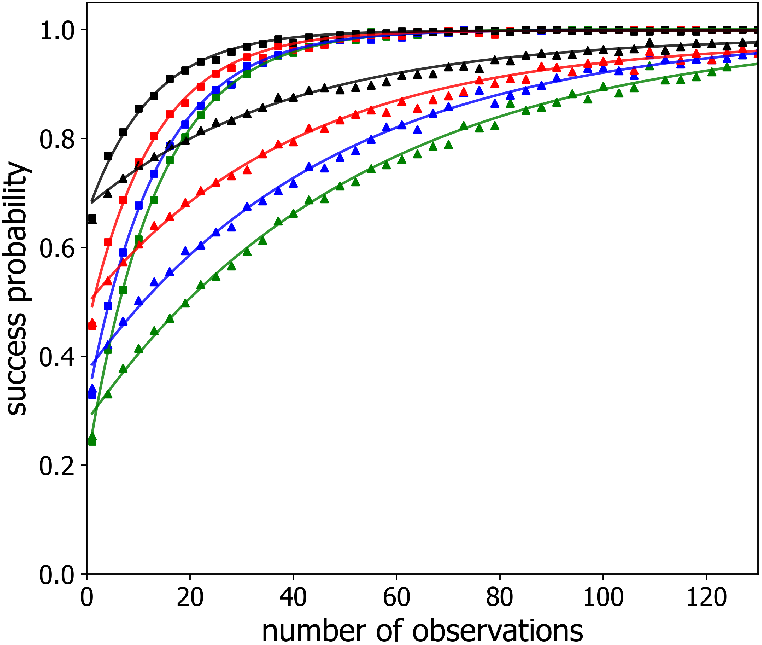
Success probability *P* (*m*) versus the number of observations *m* with an observation time step of *τ* = 1 (Eq. 1) and locus-to-locus separation of 3 (*squares*) and 6 (*triangles*) beads for varied numbers of labeled loci *n*: *n* = 3 (*black*), *n* = 4 (*red*), *n* = 5 (*blue*), *n* = 6 (*green*). a time step of *τ* = 1 (Eq. 1). For clarity, every third data point is plotted. The solid lines are single exponential fits (Eq. 9) to the data.

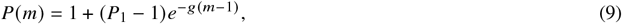

which well describes the data. The fitting parameter, *g*, is the characteristic rate at which success probability approaches unity. (The units of *g* are “per time step”, *i*.*e. g* is dimensionless.) As also can be seen from this figure, loci separated by smaller genomic distances are more rapidly mapped accurately than those separated by larger genomic distances. We ascribe this result to larger scale polymer configurational rearrangements being slower than smaller ones.

Similarly, Fig. 4 displays the success probability *P* (*m*) for a polymer with *n* = 3, 4, 5, and 6 labeled beads (black, red, blue and green triangles, respectively) separated in the chain by 4 unlabeled beads for successive observations separated in time by two different time steps, namely *τ* = 1 and *τ* = 1000 (triangles and squares, respectively). The solid lines in this figure also show the best-fit to Eq. 9. Notably, observations, separated by longer time steps, boosts the success probability, which shows a faster rate of exponential approach to the asymptote of 1 for *τ* = 1000 than for *τ* = 1. We believe this behavior is because for progressively larger values of the time step, *τ*, the polymer configurations sampled become progressively more independent of each other.

Inspection of both Fig. 3 and Fig. 4 suggests that the differences between curves for different numbers of labeled loci largely originates from their significantly different values of *P*_1_, and not from significant differences in their convergence rates. Thus, these data suggest that *g* depends only weakly on the number of labeled loci for a given time step and a given labeled loci separation. To explore this behavior in more detail, Fig. 5 plots *g* as a function of the sampling time step, *τ*, on semi-logarithmic axes, for different numbers of labeled loci, *n* = 3, 4, 7 (shown as circles, squares, triangles, respectively) and for several inter-loci spacings with 1, 3, 5, 7, 10 unlabeled beads in between (shown in blue, green, red, purple and brown points, respectively). For each combination of labeled loci number and spacing, *g* displays an S-shaped dependence on sampling time step, *τ*, with an initial increase from zero, an inflection point, and eventually a plateau value.

**Figure 5.**
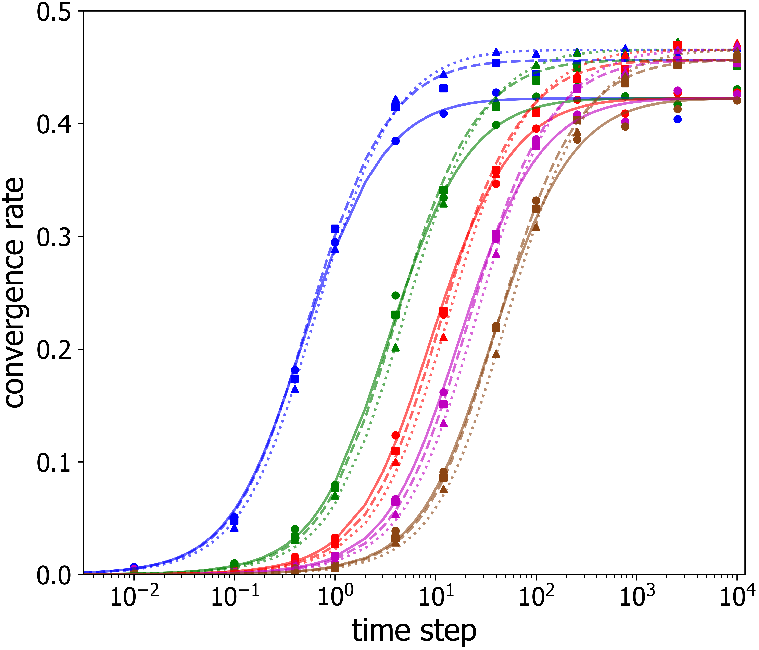
Convergence rate *g* versus timestep *τ* between successive observations for selected numbers of labeled loci *n, n* = 3 (*circles, solid lines*), *n* = 4 (*squares, dashed lines*), *n* = 7 (*triangles, dotted lines*), and varied locus-to-locus separations of 1 (*blue*), 3 (*green*), 5 (*red*), 7 (*purple*), and 10 (*brown*). The lines are the best fits of Eq. 9 to the data.

It is clear from Fig. 5 that the shapes of the *g* (*τ*) curves versus *τ* appear similar for different numbers of labeled loci and different labeled loci spacings. To quantify *g* (*τ*) and test whether it is possible to re-scale the axes, so as to collapse the *g* (*τ*) -curves for different labeled loci numbers and spacings onto a single universal curve, we employed symbolic regression to find the following empirical functional form:

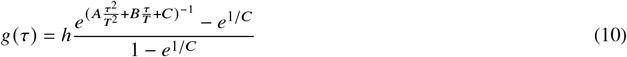

and fit it to the experimental data with *h, A, B, C* and *T* as variable parameters. Devised to mimic the measured values of *g*, Eq. 10 tends to 0 for small 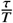, while it approaches a plateau value of *h* for large values of 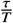. Varying the value of the “rescaling time”, *T* , shifts the function along the sampling time step axis, *i*.*e*. varying *T* varies the curve’s inflection point. First, we fit Eq. 10 to the values of *g* measured for 3 labeled loci with a spacing of 2 by varying *h, A, B, C* and *T* . Next, we fit all other *g* (*τ*) curves, as follows. For curves corresponding to three labeled loci, *h, A, B*, and *C* values were fixed to the values determined from the fit to the data for a spacing of two, while *T* only was varied to achieve a fit. For numbers of labeled loci different than three, *A, B*, and *C* were held constant, and the plateau rate, *h*, was determined from the best fit for a labeled loci spacing of 2. Then, for all other spacings for the same number of labeled loci, *h* was fixed, while *T* was varied to achieve the best fit. The resultant best-fit curves are shown as the lines in Fig. 5, demonstrating that Eq. 10 well describes the observed convergence rates. The corresponding best-fit parameter values are given in Table. 1.

We interpret the parameters, *T* and *h*, as follows. The rescaling time, *T* , which determines each curve’s inflection point, is a measure of how long it takes for one labeled polymer configuration to turn over into an independent labeled polymer configuration: for *τ* ≫ *T* , successively sampled configurations are uncorrelated. The plateau value of *g* (*τ*) , *h*, is an inverse measure of the number of observations of these uncorrelated configurations that are required to correctly identify labeled loci (Eq. 9).

Shown as the red, blue and green circles in Fig. 6 are the rescaling times, *T* , versus labeled loci spacing for 3, 4, and 7 labeled loci. For each number of labeled loci, we see that the rescaling time *T* varies relatively weakly with number of label loci, but more strongly with labeled loci spacing. In fact, the dependence of *T* on loci spacing closely follows a quadratic dependence, as shown as the solid lines in Fig. 6, which are best-fits of a quadratic function. The increase of *T* with labeled loci spacing is consistent with the expectation that larger-scale configurational changes should take longer than smaller-scale changes.

**Figure 6.**
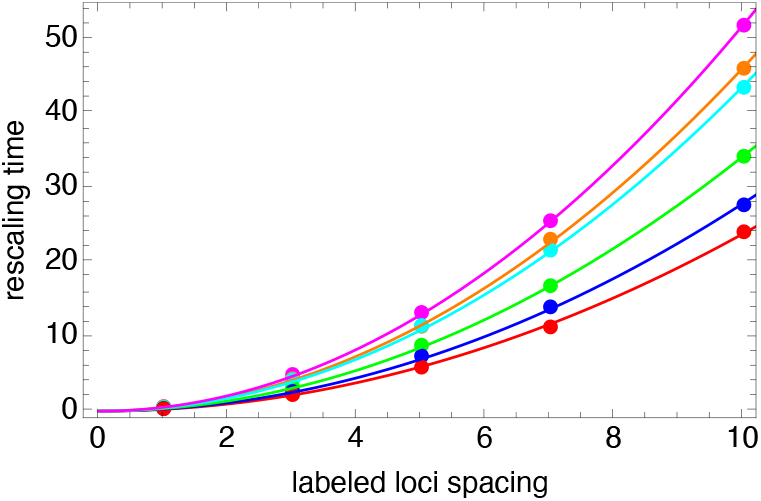
Rescaling time, *T* , versus labeled loci spacing for similarly-labeled loci and *n* = 3 (*red*), *n* = 4 (*blue*), *n* = 7 (*green*), and for alternatingly-labeled loci and *n* = 4 (*orange*), *n* = 5 (*cyan*), *n* = 7 (*magenta*). Lines are the best fit of a quadratic function of the labeled loci spacing.

Fig. 7 shows, as the green points, the best-fit values of *h* versus number of labeled loci. Importantly for the task of identifying/mapping loci, it is clear from this figure, that *h* depends only weakly on the number of labeled loci, implying that the number of independent chromatin configuration samplings required to map loci to their genomic positions correctly (i.e. with probability *P* ≈ 1) can be quite modest, even in the case of large numbers of labeled loci, when the success of correct mapping from a single observation is small (*P*_1_ ≃ 0). Indeed, for *τ* ≫ *T* , so that *g* ≃ *h*, inverting Eq. 9 gives

**Figure 7.**
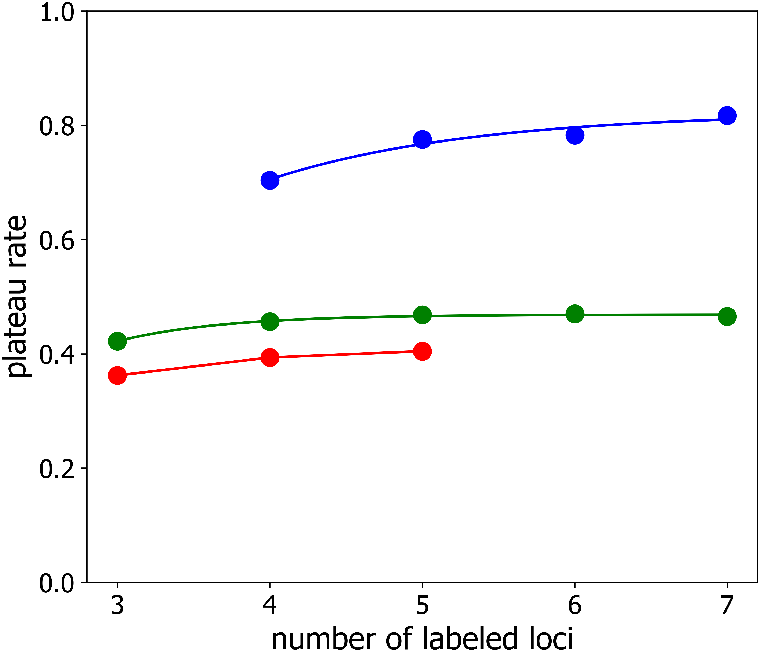
Plateau value of the convergence rate *g* versus number of labeled loci *n* for similarly-labeled loci (*green*), for alternatingly-labeled loci (*blue*), and for similarly-labeled loci on a polymer with loops (*red*). Lines are guides-to-the-eye.

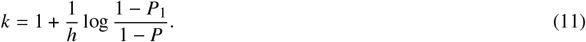

According to Eq. 11, to achieve a success probability of *P* = 0.95 even when *P*_1_ ≃ 0, the corresponding number of independent chromatin samplings is 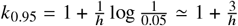. For *h* = 0.46, *k*_0.95_ ≃ 7.5, consistent with what we see in Fig. 3 and Fig. 4. Of course, for larger values of *P*_1_, reaching *P* = 0.95 will require fewer independent configuration samplings. Fig. 8 plots the normalized convergence rate versus rescaled time step, demonstrating that the data of Fig. 5 indeed collapse onto a single curve, thus confirming that normalized convergence rate, 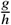, is well-described as a function of rescaled time step, 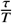, alone, i.e. 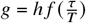.

**Figure 8.**
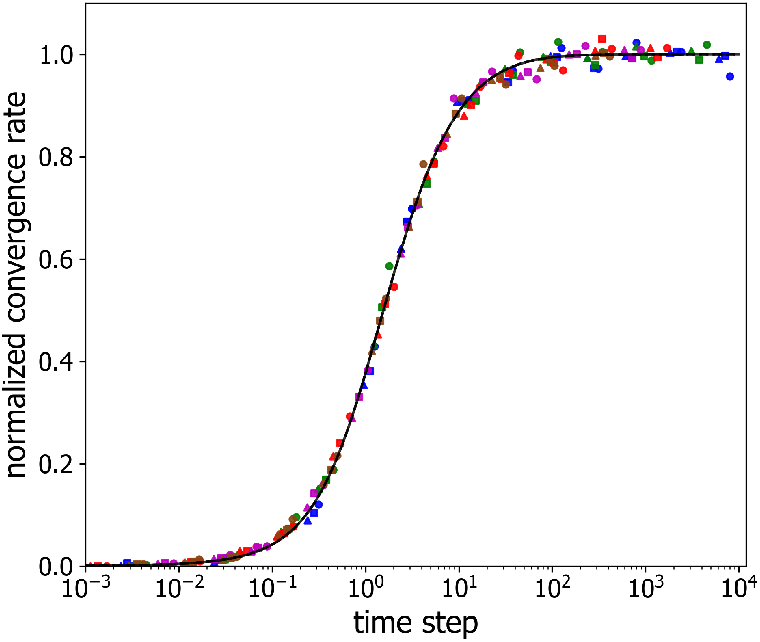
Normalized convergence rate versus rescaled observation time step. The color assignments are identical to those in Fig. 5. The solid line corresponds to the scaled version of Eq. 10.

### Success probability for alternatingly-labeled loci

Although studies visualizing loci with multiple distinguishable labels are challenging, labeling loci with two colors is feasible. For example, the two unrelated operator-repressor systems (e.g. *lacO*/LacI-GFP and *tetO*/TetR-RFP) have previously been employed together in a number of two-color studies (61–64). Alternatively, tandem arrays with two different numbers of binding sites of the same operator could be used for an approach based on two different intensities. Accordingly, we sought to investigate the case where multiple loci are visualized with two different labels, that alternate along the chromatin polymer. Because such approach provides an additional constraint on how observed loci are connected by the chromatin polymer, we reasoned that we should accordingly expect an enhanced probability of correctly mapping loci to genomic positions.

Fig. 9 shows the probability, *P*(*m*), that our procedure correctly maps observed loci as a function of the number of observations *m* for four, five, and seven loci, labeled with alternating colors/intensities with an inter-loci spacing of 4 for two different values of the time step between successive configuration samplings, namely *τ* = 1 and *τ* = 1000. As in the identically-labeled scenario, *P* (*m*) starts from its single-observation value, *P*_1_, and then increases towards unity with increasing *m*. As before, we quantified how the success probability *P* (*m*) approaches unity by fitting Eq. 9 to the data.

**Figure 9.**
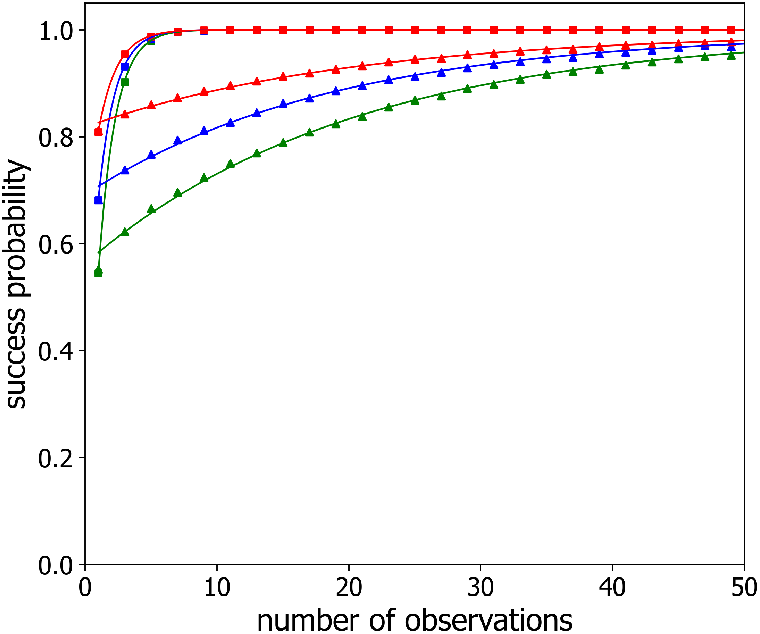
Success probability *P* (*m*) for alternatingly-labeled loci, with locus-to-locus separation of four beads, versus the number of observations *m* for observation time steps *τ* = 1 (*triangles*) and *τ* = 1000 (*squares*) and different numbers of labeled loci *n*: *n* = 4 (*red*), *n* = 5 (*blue*), *n* = 7 (*green*). The lines are best fit of single exponential (Eq. 7) to the data.

Fig. 10 displays the best-fit value of the rate, *g* (Eq. 9), for alternatingly labeled loci as a function of sampling time step, *τ*, using semi-logarithmic axes for different numbers of labeled loci and inter-loci spacings. As in the similarly labeled case, each combination of labeled loci number and spacing exhibits an S-shaped curve for the convergence rate *g* versus sampling time *τ*, where *g* (*τ*) depends relatively weakly on the number of labeled loci and relatively strongly on the labeled loci spacing. In this case too, the *g* (*τ*) curves are well described by Eq. 10, as shown by the solid lines in Fig. 10, which are the best fits to Eq. 10, following a similar procedure to that described above for the similarly labeled case. The resultant best-fit parameter values are presented in Table 1. Consequently, 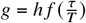 again and rescaling *g*(*τ*) curves into 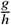 versus 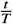 coordinates collapses all two-label data points on the same master curve, as shown in Fig. 11.

**Table 1:** Best-fit values of *h* and *T* for different numbers and different labeled loci spacing for similarly labeled and alternating-labeled loci. The values of *A, B*, and *C*, namely *A* = −0.01, *B* = −0.7663, and *C* = −0.3628, were held constant throughout.

| | $h$ | $T$ (spacing 1) | $T$ (spacing 3) | $T$ (spacing 5) | $T$ (spacing 7) | $T$ (spacing 10) |
| --- | --- | --- | --- | --- | --- | --- |
| 3 similarly labeled loci | 0.4223 | 0.3197 | 2.2203 | 5.9136 | 11.3181 | 24.1091 |
| 4 similarly labeled loci | 0.4563 | 0.3556 | 2.6406 | 7.4225 | 14.0166 | 27.7321 |
| 7 similarly labeled loci | 0.4654 | 0.4187 | 3.2216 | 8.8628 | 16.8612 | 34.2600 |
| 4 alternatingly-labeled loci | 0.7038 | 0.5755 | 4.3510 | 11.4082 | 23.1033 | 46.0845 |
| 5 alternatingly-labeled loci | 0.7753 | 0.5208 | 4.2141 | 11.5128 | 21.6406 | 43.5252 |
| 7 alternatingly-labeled loci | 0.8172 | 0.6042 | 4.9271 | 13.2642 | 25.5932 | 51.8827 |

**Figure 10.**
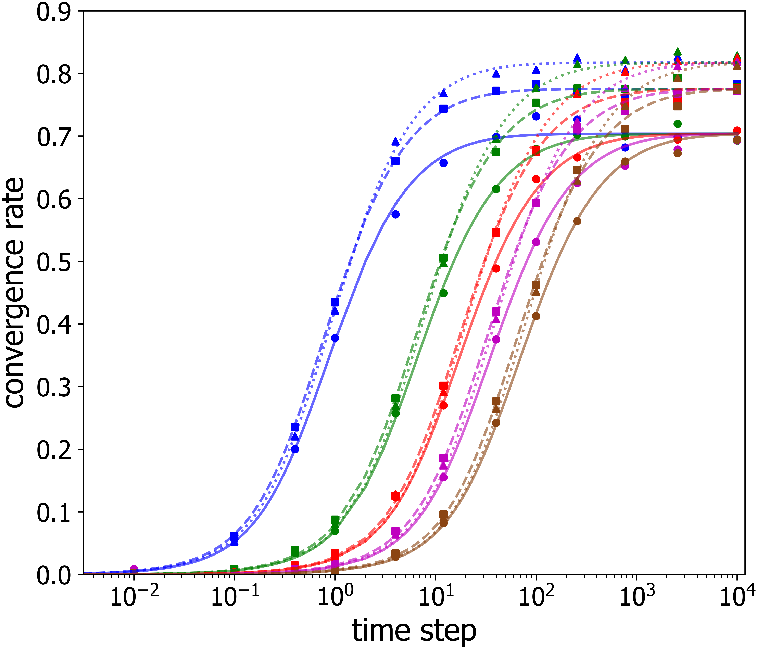
Convergence rate *g* versus timestep *τ* between successive observations for alternatingly-labeled loci. Shown are data for selected numbers of labeled loci *n, n* = 3 (*circles, solid lines*), *n* = 4 (*squares, dashed lines*), *n* = 7 (*triangles, dotted lines*), *n* = 6 (*green*), and varied locus-to-locus separations of 1 (*blue*), 3 (*green*), 5 (*red*), 7 (*purple*), and 10 (*brown*). The lines are the best fits of Eq. 9 to the data.

**Figure 11.**
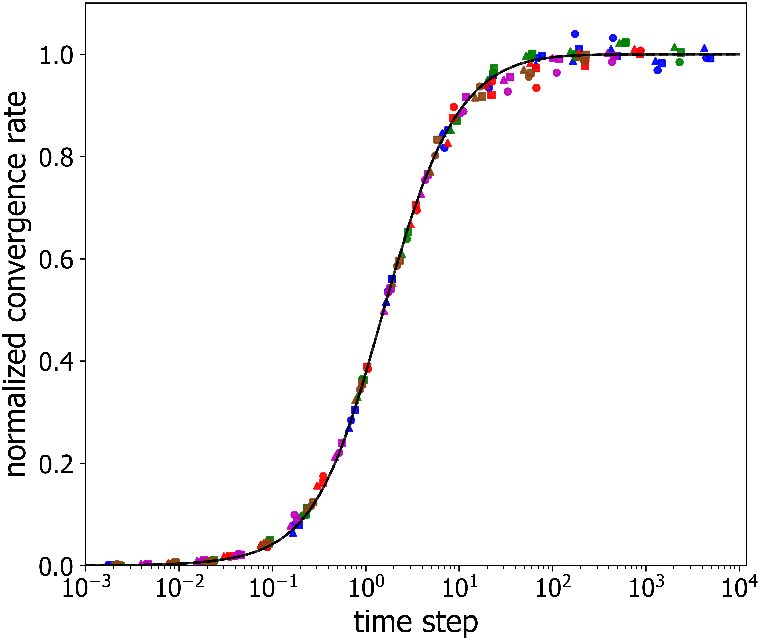
Normalized convergence rate versus rescaled observation time step in the case alternatingly-labeled loci. The color assignments are identical to those in Fig. 10. The line corresponds to the appropriately scaled version of Eq. 10.

Similarly to the same-label case, the rescaling time, *T* , varies quadratically with inter-loci separation for alternatingly labeled loci, as shown in Fig. 6 as the green, orange and magenta points and lines, while the plateau value *h* varies only slightly versus number of label loci, as shown is the blue points in Fig. 7. Importantly, although the the rescaling time, *T* , is longer for the two-label case (Fig. 6), the plateau value, *h*, is also higher (Fig. 7), implying that significantly fewer independent polymer configuration samplings are needed to determine the correct loci mapping in the case of alternatingly-labeled loci, as can be seen in Fig. 9 for the *τ* = 1000 curves.

### Success probability for similarly labeled loci on independent polymers with loops

As noted in the Introduction, chromatin in living cells realizes a dynamic configuration of loops that are inhomogeneously and stochastically distributed along the chromatin polymer. To investigate how the presence of dynamic loops might effect the probability of correct loci mapping, we carried out simulations of a modified Rouse model (35, 47, 59), that includes stochastically appearing, growing and disappearing loops. In this case, we chose to sample polymer configurations at times corresponding to *t* ≫ *T*, i.e. when subsequent observations sample independent polymer configurations.

Fig. 12 shows the success probability, *P* (*m*) , versus the number of independent observations, *m*, of the looped polymer configurations for three different numbers of labeled loci, *n* = 3, 4, 5, and three different inter-loci spacings, namely 800, 1000 and 1200, which are all much larger than the mean loop size of 30. As expected for polymer configurations, that are independent at all length scales, *P* (*m*) does not depend on the inter-loci separation in this case. As for the classical Rouse model, the success probability is well described by Eq. 9. In fact, the best-fit values of *P*_1_ are very similar to those found from simulations of the classical Rouse model without loops, as shown by the red points in Fig. 2. In addition, as shown by the red points in Fig. 7, the best-fit values of the characteristic convergence rate in this case are comparable, but slightly smaller than, the plateau rate from simulations of the classical Rouse model without loops. We ascribe this small reduction in plateau rates to the additional stochasticity, introduced by loop extrusion.

**Figure 12.**
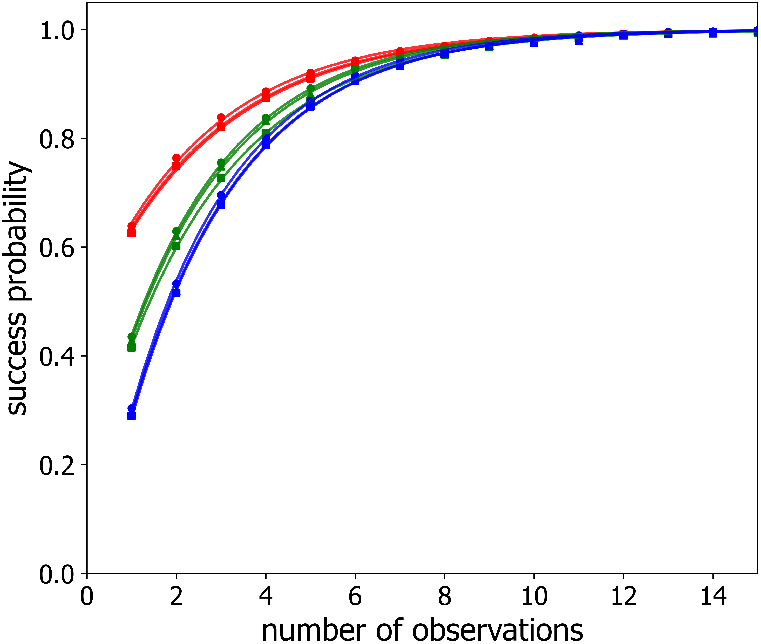
Success probability *P* (*m*) versus the number of observations of independent polymer configurations *m* for a Rouse polymer with loops. Shown are data for selected numbers of labeled loci *n, n* = 3 (*red*), *n* = 4 (*green*), *n* = 5 (*blue*) and varied locus-to-locus separations of 800 (*squares*), 1000 (*triangles*), 1200 (*circles*) beads. The lines are the best fits of Eq. 9 to the data.

## DISCUSSION

In spite of recent success in elucidating its spatial organization, chromatin’s behavior along the fourth dimension, time, remains far less studied, and correspondingly much less understood. To address the critical need to elucidate chromatin dynamics, this paper proffers a route towards an experimental characterization of coarse-grained chromosomal dynamics, via particle tracking of multiple labeled loci, labeled with one or two different fluophor colors or intensities. Building on the hypothesis that single-particle-tracking measurements should enable determination of the correct order along chromatin of similarly labeled loci because proximity along the genome leads on-average to physical proximity, we have found that the probability of correctly assigning loci, on the basis of minimizing loci square separations, approaches one exponentially as the number of observations increases. Remarkably, the characteristic convergence rate at which the probability approaches unity with increasing numbers of samples depends only weakly on the number of labeled loci. This implies that even for a large number of identically-labelled loci, when the number of possible mappings grows factorially, correct mapping can be identified with high fidelity by tracking them across only about 8 independent chromatin configurations.

In the case of multiple loci labeled with two distinguishable labels (such as fluorophors with different colors), that alternate along the chromatin polymer, we find that the convergence rate takes larger values than for similarly labeled loci, meaning that about 5 independent chromatin configurations are needed to establish loci identities. For an improved model of chromatin, namely a modified Rouse-model polymer that realizes a population of dynamic loops, we find that for independent looped polymer configurations, the success probability approaches unity with increasing numbers of polymer configurations in a similar fashion to that found for independent configurations of the classical Rouse model polymer.

These collected results suggest that particle tracking of multiple labeled loci over time may be a feasible way to establish experimentally the coarse-grained configuration of the chromatin polymer in individual living cells. A key question, though, is how does the time for chromatin to proceed through the required number of independent configurations compare to the hard-limit on the duration of a single-particle tracking experiment set by fluorophor photo-bleaching, that occurs as a result of collecting images sufficiently closely in time to achieve faithful tracking (55). An important clue concerning the relevant time scales comes from extant single particle tracking experiments, such as Refs. (43, 44), which measured the mean square displacements (MSDs) of individual labeled loci versus time in budding yeast. Typically, in these experiments, with increasing time, the MSDs increase from a small value at few-second time scales to achieve a plateau value at about 20 or 30 s, which is then maintained for the total duration of the experiment (up to 600 s). The observation of a plateau value implies that each gene locus fully explores the volume available to it on on the 20-30 s time scale. Since the chromatin polymer is comprised of multiple such loci, it seems plausible, although not proven, that the chromatin configurational rearrangement time is on the order of 20-30 s, at least in budding yeast. Since this time scale is significantly shorter than the duration of the experiments in Refs. (43, 44), we are optimistic that single-particle-tracking of multiple labeled loci will enable determination of their correct order along chromatin. Importantly, the large number of labeled loci does not pose a real challenge in assigning loci identity in this approach. Instead, the true limitation to the granularity of the chromatin representation depends on the number of loci that can be accurately detected and tracked.

## 1 AUTHOR CONTRIBUTIONS

A. M., A.I.R., T. Y. I. S. M. C. K. and S. G. J. M. concieved the research and wrote the paper. A. M., T. Y. and S. G. J. M. carried out simulations. A. M. and S. G. J. M. analyzed the simulation data.

## ACKNOWLEDGEMENTS

This research was supported by NSF EFRI CEE Award EFMA-1830904 and NSF Physics of Living Systems via Award 2412859. A. I. R. was supported by NIH Grant T32GM145469.

## DECALREATION OF INTERESTS

The authors declare no competing interests.

